# Individual vulnerability to stress is associated with increased demand for intravenous heroin self-administration in rats

**DOI:** 10.1101/627471

**Authors:** Nathaniel P. Stafford, Theodore N. Kazan, Colleen M. Donovan, Erin E. Hart, Robert C. Drugan, Sergios Charntikov

**Affiliations:** Department of Psychology, University of New Hampshire

**Keywords:** heroin, stress, swim-stress, post-traumatic stress disorder, heroin demand, economic demand, stress reinstatement, cue reinstatement

## Abstract

Opioid use is a widespread epidemic, and traumatic stress exposure is a critical risk factor in opioid use and relapse. There is a significant gap in our understanding of how stress contributes to heroin use, and there are limited studies investigating individual differences underlying stress reactivity and subsequent stress-induced heroin self-administration. We hypothesized that greater individual vulnerability to stress would predict higher demand for heroin self-administration in a within-subjects rodent model of stress and heroin use comorbidity. Male rats were exposed to inescapable intermittent swim stress and individual biological (corticosterone) or behavioral (open field, social exploration, and forced swim tests) measures were assessed before and after the stress episode. Individual demand for self-administered heroin (0.05 mg/kg/infusion; 12-hour sessions) was assessed using a behavioral economics approach followed by extinction and reinstatement tests triggered by stress re-exposure, non-contingent cue presentations, and yohimbine (0, 1.0, or 2.5 mg/kg). We found that behavioral, biological, and a combination of behavioral and biological markers sampled prior and after the stress episode that occurred weeks before the access to heroin self-administration predicted the magnitude of individual demand for heroin. Non-contingent presentation of cues, that were previously associated with heroin, reinstated heroin seeking in extinction. For the first time, we show that individual biological response to an ecologically relevant stressor in combination with associated behavioral markers can be used to predict subsequent economic demand for heroin.

## 1 Introduction

People subjected to traumatic experiences are especially susceptible to the abuse of depressants, including opioids, but there is evidence of individual vulnerability or resilience to such stresses. Traumatic psychological stress, often accompanied by physical trauma, can include harassment, sexual abuse, bullying, domestic violence, life-threatening events, and other individually subjective negative experiences. It is known that stress is an important factor in the development of addiction disorders, and it is generally understood what critical neurobiological substrates underlie this mechanism (Sofuoglu et al. 2014). What is not known is how individual vulnerability or resilience to stress relates to susceptibility to use and abuse of opioids. Heroin use in the US is on the rise and without effective treatment options to combat this change (Jiang et al. 2017). Understanding individual underlying factors linking stress vulnerability to opioid use may lead to the development of more efficacious individualized treatment approaches than what is currently available.

Substance use and addiction are strongly associated with risk factors like stress and mental trauma (Perkonigg et al. 2000; Breslau et al. 2003; Zywiak et al. 2003; Dewart et al. 2006; Mills et al. 2006). The evidence for stress and heroin use comorbidity is less extensive as for other drugs of abuse, but the existing findings from epidemiological research, and few preclinical studies, strongly link stress with opioid use. For example, in one epidemiological study (Cottler et al. 1992) 43% of polydrug or cocaine/opiate users reported experiencing traumatic stress in the past and met DSM-III criteria for posttraumatic stress disorder (PTSD; odds ratio = 5.06; nearly twice the rate for other drugs). In another epidemiological study, 88% of people that abused opioids had been exposed to traumatic stress and the highest prevalence of PTSD was among individuals with an opioid use disorder (Mills et al. 2006). Importantly, these studies show that not every individual with such adverse experiences transitions to abuse opioids.

There is limited evidence from preclinical studies implicating stress in increased opioid self-administration. For example, rodent studies show that daily immobilization stress increased oral consumption of both heroin and fentanyl (Shaham 1993), while daily footshock stress increased lever-press responding for liquid fentanyl in an operant self-administration paradigm (Shaham et al. 1992). Subsequent work by Shaham and Stewart (1994) demonstrated that intermittent footshock stress also increased intravenous self-administration of heroin. The study by Shaham and Stewart (1994) provided direct evidence of stress-induced enhancement of heroin reinforcement obtained using a preclinical model of intravenous self-administration. Additional preclinical studies also found that physical and pharmacological (yohimbine) stress can also reinstate heroin-seeking behavior in extinction (Shaham and Stewart 1995; Shaham et al. 1996; Banna et al. 2010). Furthermore, there is a number of studies that have investigated the role of individual differences in preclinical animal models of addiction (Deroche-Gamonet et al. 2004; McNamara et al. 2010; Belin and Deroche-Gamonet 2012; Dilleen et al. 2012; Koffarnus and Woods 2013; for review see Belin et al. 2016). For example, Deroche-Gamonet et al. (2004) showed that only a small percentage of rats show addiction-like behaviors when assessed using extended cocaine self-administration paradigm, while Dilleen et al. (2011) showed that only high-anxious rats exhibited a higher pattern of escalation of cocaine but not heroin self-administration in comparison to less anxious rats. Although there is a strong body of preclinical evidence demonstrating that stress can contribute to heroin taking or seeking behaviors and that these effects may vary on the individual level, there is a significant gap in our understanding of how individual sensitivity to stress interacts with these behaviors.

Many individuals are exposed to traumatic stress in their lifetime; however, only a small proportion eventually transition to use and abuse opioids (Cottler et al. 1992; Breslau et al. 2003; Dewart et al. 2006; Mills et al. 2006). Based on this premise, we conceptualized a hypothetical human model where vulnerability to stress mediates stress-induced progression to heroin use and abuse. In that theoretical model, many individuals are exposed to traumatic stress, but only those that are vulnerable to traumatic stress have a higher risk of using or abusing heroin after a traumatic stress experience. In humans, vulnerability to traumatic stress is often defined as exhibiting long-lasting symptoms after the stress episode and may include hyperarousal, hypervigilance, social withdrawal, and cognitive alterations, to name a few. We further hypothesized that we could simulate some of these individual effects in a preclinical animal model of stress and heroin use comorbidity. In that preclinical rodent model of stress and heroin use comorbidity, all rats are first exposed to a stress episode and then, after a period of time that allows for the development of long-term stress effects, assessed for heroin consumption using behavioral economics approach. Importantly, rather than investigating the effects of stress on rates of heroin self-administration, the approach that often requires additional non-stress controls, this model uses a within-subjects design and focuses on understanding how individual variability in reactivity to stress relates to heroin taking. Thus, in this preclinical model of stress and heroin use comorbidity, all rats are exposed to a stressor, and all subsequent individual effects are assessed as they unfold over time using mixed-effects linear modeling that is especially suitable for these types of designs (Glass and Mackey 1988; O’Connor 1990).

In the present study, we used intermittent swim stress (ISS) to induce long-lasting effects of stress. The ISS protocol is derived from both the learned helplessness and forced swim test (FST) models (Brown et al. 2001; Drugan et al. 2013). ISS is an effective stressor that induces long-lasting behavioral symptoms analogous to cognitive deficits (Christianson and Drugan 2005; Levay et al. 2006; Drugan et al. 2009, 2014), behavioral despair (Christianson and Drugan 2005; Drugan et al. 2014), social anxiety (Stafford et al. 2015), altered drug reactivity (Brown et al. 2001; Drugan et al. 2007), and is sensitive to pharmacological treatments like selective norepinephrine reuptake inhibitors (Drugan et al. 2010; Warner and Drugan 2012). It is important to note that cold water is a natural stressor for a rat that they can encounter in the environment and thus it is a stressor with high ethological relevance inducing relevant behavioral and neurobiological responses (for review see Drugan et al. 2016). Individual reactivity to ISS-induced stress episode can be assessed by measuring biological or behavioral stress that can be then correlated with subsequent economic demand for heroin. We hypothesized that rats that are vulnerable to the effects of stress would have long-lasting effects that will translate to higher demand for heroin long after the stress exposure. We here show that a) rats vary in their biological response to stress and the behavioral responses sampled before and after the stress episode, b) that variation can be conceptualized as a continuous phenotype ranging from vulnerability to resilience, and c) vulnerability to stress, in this model, predicts higher economic demand for self-administered heroin.

## 2 Materials

### 2.1 Animals

A total of 24 male adult (PD 70-90) and 12 juvenile (PD 28-32) Sprague Dawley rats (SAS Derived, Charles River Labs, Kingston, NY, USA) were used in the study. Juvenile Sprague Dawley males served as social exploration stimuli in the social exploration tests. Upon arrival at the vivarium, adults were single housed and acclimated to a colony for at least one week prior to experimentation. The vivarium was maintained on a 12 h light/dark cycle with lights on at 0700. For all rats, food and water were available ad libitum. All procedures were in accordance with the Guide for the Care and Use of Laboratory Animals, Eighth Edition (Institute for Laboratory Animal Research, The National Academies Press, Washington, DC, 2011) and were reviewed and approved by the University of New Hampshire Institutional Animal Care and Use Committee.

### 2.2 Apparatus

#### 2.2.1 Social Exploration

Social exploration pretest and posttest were conducted in identical test chambers, which consisted of a plastic tub cage (40.6 cm×20.3 cm×20.3 cm; l×w×h), wire lid, and 3 cm of wood shaving bedding free of food and water. The room was lit by cool fluorescent bulbs and light penetration into the test chamber was 200-300 lx. Juvenile stimuli were used for a maximum of 4 tests and adults were never exposed to the same juvenile more than once. A camera that was mounted above the apparatus recorded behavior during each test session (Stafford et al. 2015).

#### 2.2.2 Open Field

Open-field tests were conducted in an open-top square plywood box (120 cm×120 cm×25 cm; l×w×h) painted with flat black enamel. Test sessions were video recorded from a camera mounted above the apparatus and processed using Ethovision XT software (version 8.5; Noldus Information Technology, Wageningen, The Netherlands).

#### 2.2.3 Intermittent Swim Stress

Intermittent swim stress was conducted in two acrylic cylinders (21 cm×42 cm; d×h) with a 6.35 mm galvanized wire mesh at the bottom of each cylinder. Cylinders were suspended over a tank (80.6 cm×45.7×28.6 cm; l×w×h) filled with water maintained at 15±1 °C. The apparatus was controlled by a Med-PC interface and software (Med Associates, Inc.; St. Albans, VT, USA). Space heaters, above and in front of each cylinder, circulated warm air (~36 °C) in and around the cylinders to limit the effects of hypothermia during the inter-trial intervals.

#### 2.2.4 Forced Swim Test

The forced swim test was conducted in acrylic cylinders (20 cm×46 cm; d×h). The water was filled to 30 cm height and was kept at 24 °C. Test sessions were video recorded and quantified as described above.

#### 2.2.5 Self-administration chambers

Self-administration chambers (ENV-008CT; Med Associates, Inc.; St. Albans, VT, USA) measuring 30cm×20 cm×20 cm (l×w×h), were enclosed in a sound- and light-attenuating cubicle equipped with an exhaust fan. Each chamber was equipped with two retractable levers, two stimulus lights, and a house light. The infusion pump, that was located outside of the sound-attenuating cubicle, was equipped with a syringe that was connected to a swivel inside the chamber and that was further extended through a spring leash suspended over the ceiling of the chamber on a balanced metal arm. Lighting in the test room was maintained at 400-500 lx and exposure to light was minimized during the transfer from the colony.

### 2.3 Drugs

Diamorphine hydrochloride (generously provided by NIDA Drug Supply Program) and yohimbine hydrochloride (Sigma; St, Louis, MO, USA) were mixed in a 0.9% sterile saline solution.

## 3 Methods

Experimental progression is shown in Figure 1. At the start of experimentation, all rats received twice daily handling for one week by all experimenters with the last three days including towel restraint habituation for blood collection. Rats were first subjected to a series of pretests consisting of social exploration and open field. On the following day, rats were subjected to the intermittent swim stress, an ethologically relevant stressor with high ecological validity (Brown et al. 2001; Drugan et al. 2005, 2010, 2016). Twenty-four hours after the intermittent swim stress, rats were subjected to a series of post-tests that included social exploration, open field, and FST. In order to assess stress-induced changes in behavior and corticosterone, stress reactivity was tested during the light phase when basal glucocorticoid levels nadir. Eighteen to twenty-two days after the initial ISS-induced stress episode rats started heroin self-administration that consisted of multiple phases including the acquisition of self-administration, assessment of individual demand for heroin, and reacquisition of self-administration. After that, rats underwent extinction and reinstatement testing with heroin no longer available.

**Figure 1.**
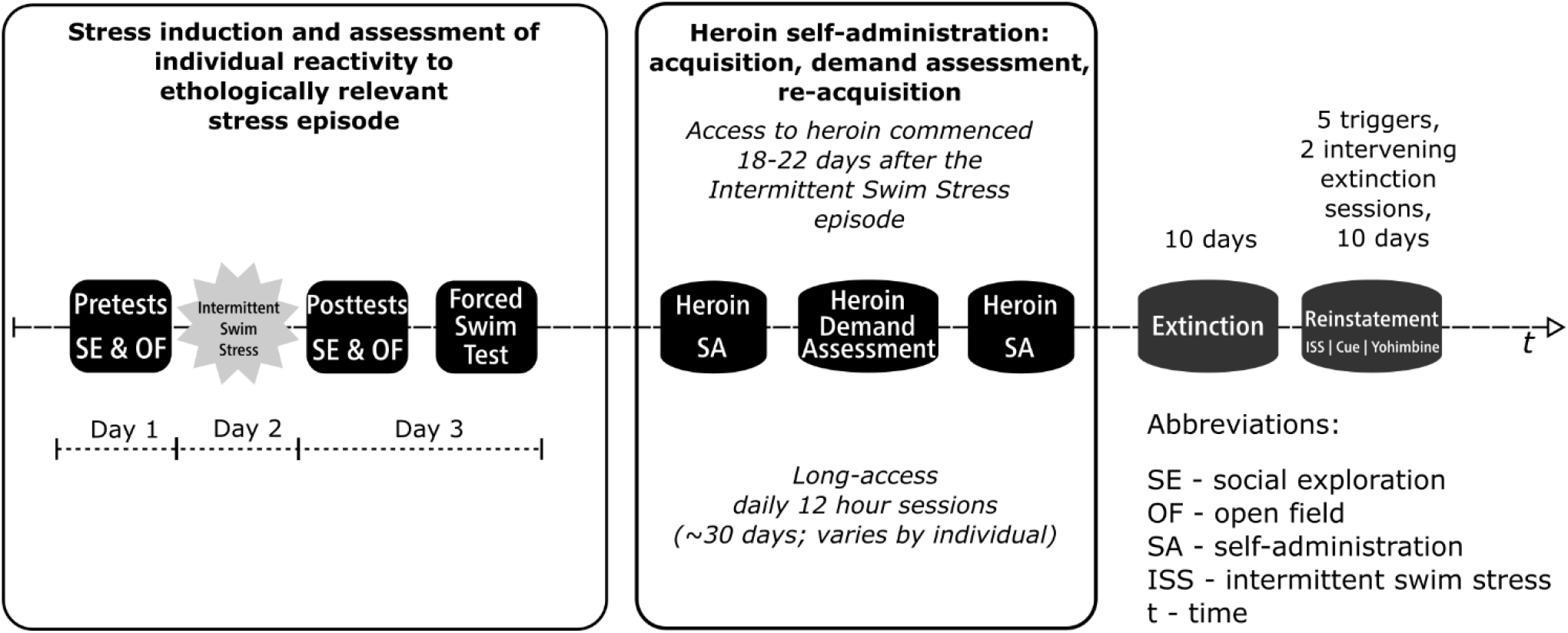
Experimental progression. The study used a within-subjects design (final N=16; 8 rats removed from the study due to patency loss) with several distinct experimental phases occurring sequentially as outlined in the figure. <u>The first phase modeled exposure to a stress episode</u>. During that phase, rats were subjected to Intermittent Swim Stress and their individual reactivity to stress was assessed using a series of pretests and posttests that included social exploration, open field, and forced swim test (see first 4 blocks of the timeline). <u>The second phase modeled drug taking</u> (see Heroin SA, Heroin Demand Assessment, and Heroin SA blocks). In the second phase, rats were allowed to self-administer heroin (0.05 mg/kg/infusion; 12 hours/day) and their demand for heroin was assessed using behavioral economics model; rats were trained to self-administer heroin before this assessment and retrained after to ensure a stable level of responding. <u>The third phase modeled abstinence</u> (see Extinction block). Extinction phase was identical to self-administration phase except that heroin and cues that were previously paired with heroin infusions were no longer available. <u>Fourth and final phase modeled relapse</u> (see Reinstatement block). In this final phase, the resurgence of active lever responding in extinction was triggered by an abbreviated version of the intermittent swim stress, pharmacological stressor yohimbine (0, 1.25, and 2.5 mg/kg), or a non-contingent cue presentation.

### 3.1.1 Social Exploration

Social exploration consisted of a 1 h acclimation to the test chamber after which a juvenile was placed into the cage for 3 min. Social exploration tests were analyzed in real time by two experimenters (interrater reliability: r = 0.93). The following behaviors of the adult directed toward the juvenile were quantified: sniffing (direct snout contact against any portion of the juvenile and primarily directed at the anogenital region), pinning (minimum of two fore-paws against juvenile), allogrooming, and chasing. A total sum of these measures comprised the reported social exploration score.

### 3.1.2 Open Field

Rats were placed individually into the center of the open field apparatus for 10 min, after which they were returned to the vivarium. Locomotor activity, defined as the distance traveled (path length), and time spent in the center vs. perimeter (thigmotaxis) were measured using Ethovision XT 8.5. Ethovision software was calibrated to identify and track the center point of the white subject against the black arena background using the static subtraction method (subject contour detection was set to erode first, then dilate, at 1 pixel for optimal subject detection) at the default 5 samples/s for the Open Field Template within the program. The following parameters were defined in Ethovision for the locomotor variables. The open-field apparatus was divided into two portions: the center consisted of a central 60cm × 60cm square (located 30 cm from the apparatus wall), while the remaining surrounding area of the apparatus consisted of the perimeter. The path length was calculated as total distance moved (in cm) of the center point of the subject throughout the entire arena and movement was defined when the center point changed position above a threshold of 1 cm/s. All dependent measures were divided into the first 5 min (habituation; 0-5 min) and last 5 min (test; 5-10 min) of the test. Behaviors during the second 5 min bin were used for data analyses.

### 3.1.3 Exposure to a stress episode: intermittent stress swim

ISS was administered 24 hours after pretests between 0700-1230. Each swim trial consisted of a five-second forced swim in which the cylinder was submerged to a depth of 25 cm. One hundred trials were presented at a variable 60 s (10-110 s) inter-trial-interval. Immediately following ISS, rats were towel-dried and returned to the vivarium to heated cages (heating pad placed underneath cages) to promote vasodilation for blood collection that occurred thirty minutes post-ISS and was performed in a separate room.

### 3.1.4 Forced Swim Test

The FST was administered immediately following the post-ISS social exploration and open field tests. During the FST, rats were forced to swim for 5 min in 24±1 °C. Water depth was kept at 30 cm. This depth level forced rats to swim because it avoids the possibility of body support by the tail touching the bottom of the cylinder (i.e., tail-standing). Forced swim test videos were analyzed via a widely used and reliable serial time-sampling procedure in which behavior is scored as either immobile, swim or climb in 5 s intervals and reported as mean counts for each behavior (Detke et al. 1995; Drugan et al. 2010). Immobility was defined as only necessary movements to keep the head above water. Swimming was defined as active movements, including dives that did not involve struggling against the side of the cylinder. Climbing was defined as active struggling against the side wall of the cylinder in which the fore-paws broke the surface of the water. Inter-rater reliability was calculated for all measures (immobility, *r*= 0.96; climbing, *r*= 0.97; swimming, *r* = 0.94).

### 3.1.5 Blood Collection and analysis

Twenty-four hours before initiation of behavioral tests, a baseline blood sample was collected from the lateral tail vein; all samples were collected between 0900-1300. Second blood collection occurred 30 min after ISS test. Additional blood collections were performed 30 min post-FST when circulating corticosterone concentration peaks (Connor et al. 1997), the day after the last self-administration session, and 30 min after the abbreviated ISS exposure that was used as one of several reinstatement triggers. Rats were lightly restrained in a towel; the tail was placed into 46±2°C water to promote vasodilation and approximately 300 microliters blood was collected via lateral tail vein incision with a #11 scalpel into a capillary tube. The first incision was made in the distal 2 cm of the tail with subsequent incisions made at least 1 cm rostral of the previous. All samples were collected within 3 min and rats were returned to a home cage within 5 min (Fluttert et al. 2000; Drugan et al. 2005). Samples were centrifuged at 4°C for 4 min at 1300 rpm to separate red blood cells and extract plasma, which was stored at −80°C until assay. Samples were analyzed via enzyme-linked immunoabsorbent assay (Arbor Assays, Ann Arbor, MI, USA) and processed in duplicates. Corticosterone concentrations were read at 405 nm on a BioTek microplate reader using Gen5 software (BioTek, Winooski, VT, USA). The intra-assay and inter-assay coefficients were 6% and 8%, respectively.

### 3.1.6 Catheter Implantation Surgery

Subjects were anesthetized with 1 ml/kg ketamine (100 mg/ml) and xylazine (20 mg/ml) mixture (2:1 ratio; administered intramuscularly; Sigma; St. Louis, MO, USA). A polyurethane catheter (22 Ga; RJVR-23; Strategic Applications Inc.; Lake Villa, IL, USA) with a rounded tip and double suture beads (one secured internally and other externally) was implanted into the right external jugular vein. The other end of the catheter was subcutaneously placed around the shoulder and exited below the scapula via subcutaneously implanted polycarbonate back-mount access port (313-000BM; Plastics One Inc.; Roanoke, VA, USA). Immediately following the surgery, catheters were flushed with 0.2 mL mixture of 30 U/ml heparin and 50 mg/mL of cefazolin (antibiotic) diluted in sterile saline (0.9% NaCl). Atipamezole hydrochloride (0.5 mg/kg; IM; Sigma; St. Louis, MO, USA) diluted in saline was used to terminate anesthesia. To manage post-surgical pain, butorphanol tartrate (1 mg/kg; SC) was administered immediately after the surgery and daily for the next two recovery days. Starting from the day after surgery, catheters were flushed daily with 0.2 mL heparinized saline (30 U/ml) and cefazolin (50 mg/mL) mixture. Catheter patency was assessed when patency loss was suspected or upon completion of the self-administration phase with an infusion of 0.05 ml xylazine (20 mg/ml; IV). This xylazine concentration produces clear motor ataxia within 5-10 s (Charntikov et al. 2013; Pittenger et al. 2018). Rats that did not exhibit noticeable motor ataxia within 5-10 s following xylazine infusion were considered non-patent.

### 3.1.7 Preliminary training

Rats were trained to lever press over 3 daily sessions. At the start of each session, the house-light was turned on and a randomly selected lever (right or left) was inserted. A lever press or lapse of 15 s resulted in sucrose delivery (100 μl of 30% liquid solution; 4-s access) via a raised dipper, lever retraction, and commencement of a timeout (average=60 s; range=30 to 89 s). Following the timeout, a randomly selected lever was inserted with the condition that the same lever could not be presented more than twice in a row. This protocol was repeated for 60 sucrose deliveries. Sessions lasted 65 to 80 min depending on individual performance. Training continued until a lever press was made on at least 80% of the lever insertions for two consecutive days (i.e., 3 to 5 sessions). After rats met this criterion, they were surgically implanted with an intravenous catheter as described earlier. Following 7 days of recovery, lever press training continued as described above, but the response contingency was changed to a variable ratio (VR_3_) schedule of reinforcement where on average every third response was followed by a sucrose delivery (range=1 to 6 presses). At least 80% of the 60 available sucrose deliveries had to be earned to move to the self-administration phase; this occurred after 3 to 5 sessions. This protocol ensures high rates of responding, yet both levers have similar reinforcement history to avoid any potential bias of differential lever press training in later phases.

### 3.1.8 Heroin self-administration and assessment of individual demand for heroin

After recovery from surgery, all rats were retrained to lever press on both levers using a variable schedule of reinforcement (VR_3_) and after meeting a criterion transitioned to daily 12 h heroin self-administration (0.05 mg/kg/infusion; VR_3_). Each session began with a termination of the house light, insertion of both levers, and a 0.9 s infusion to flush approximately 90 % of catheter volume. Completion of the required response resulted in a ~1 s infusion of heroin, retraction of both levers, and illumination of both cue lights above each lever for a 20 s timeout. Additional 5 min timeout with levers retracted and house light illuminated was instituted every 55 min to mitigate possible overdose-related deaths. All rats self-administered the exact dose of heroin using a variation in infusion duration that was automatically controlled by the program based on their pre-session weight. To closely simulate drug taking conditions observed in humans, all self-administration sessions were conducted during the night cycle which corresponds to rodents’ active phase (1900-0700). Access to heroin self-administration commenced 18-22 days after the initial ISS-induced stress episode. After 8 days of heroin self-administration on VR_3_ as described above, heroin was earned on fixed ratio (FR) schedule of reinforcement that was escalated daily using the following sequence: 1, 3, 5, 8, 12, 18, 26, 38, 58, 86, 130, 195, 292, 438, and 657. Subsequently, rats were allowed to self-administer heroin on VR_3_ as described above for an additional 5 days.

### 3.1.9 Extinction and reinstatement

Extinction training was identical to self-administration sessions except that active lever responding had no programmed consequences; no heroin, cues, or lever retractions. Reinstatement tests commenced on the day after last (10th) extinction session at the usual time of self-administration or extinction sessions (1900). There were a total of five one-hour reinstatement tests with two intervening daily extinction sessions between these tests. Reinstatement triggers included abbreviated ISS re-exposure (20 trials), non-contingent cue presentations, and yohimbine (0, 1.0, or 2.5 mg/kg; IP). Abbreviated ISS and yohimbine were administered 30 min before the beginning of each test. Non-contingent cue presentations were identical to the cues that were associated with heroin infusions and consisted of both cue lights turned on, and both levers retracted for 20 seconds. These cue triggers were presented at the start of the session and every five minutes from the beginning of the session thereafter. The order of reinstatement tests for each rat was assigned using a Latin square design.

### 3.2 Analytical approach

Data from eight rats were removed from the study due to intravenous catheter patency loss (final N=16). Some individual corticosterone datum was not available because of failed sampling attempts or not enough serum for the analysis (two samples after the FST, one sample after extinction, and three samples after stress-induced reinstatement). Statistical analyses involving corticosterone data were restricted to subjects with available data. Individual demand for heroin was derived from the amount of heroin consumed (mg/kg) over each FR schedule of reinforcement (Hursh and Silberberg 2008). Essential value from the demand model was used to estimate individual demand for a reinforcer and was calculated from the nonlinear least squares regression model fit to the individual consumption data from each schedule of reinforcement using the following formula: *log Q* = *log Q_o_* + *k*(*e*^−*αQ_o_C*^ − 1) where *Q* represents reinforcer consumption, *Q_o_* is a consumption when price is zero or free, *κ* is a constant for the range of demand, *e* is the base of the natural logarithm, *C* is the varying cost of each reinforcer, and *α* is the rate of decline in relative log consumption with increases in price. The essential value was calculated from the demand model using the following formula: *EV=1/(100 × a × k^1.5^)*. The main advantage of using essential value is that it is a unifying measure based on several critical parameters forming exponential-demand equation. For example, essential value takes into consideration consumption when the price of a reinforcer is low (e.g., FR1), when the price of the reinforcer is high (e.g., higher or terminal FR schedules), and the slope of the demand curve - also referred to as elasticity. Using this approach that we previously demonstrated in nicotine self-administration study, it is possible to plot individual demand curves for each subject, and most importantly, derive a single value of individual demand for heroin that is based on performance over a range of schedules of reinforcement (Kazan and Charntikov 2019).

The difference between pre- and post-stress behaviors were assessed using t-tests (performed using R 3.4.2; {stats} package). Responding on active and inactive levers was analyzed using ANOVAs (performed using GraphPad Prism). Individual effects were analyzed using linear mixed-effects modeling with maximum likelihood fit (performed using R 3.4.2; {lme} package). Individual effects were analyzed by building a model with a maximum likelihood fit from a baseline that does not include any predictors other than an intercept. The model was then built by adding one predictor or a combination of predictors and comparing it to a baseline. The model fit was declared significant when its addition improved the model by accounting for significantly more variance (the fit was examined using the Likelihood Ratio test of fixed effects; p<0.05). The proportion of variance explained by the factors is reported as the marginal R^2^. Additional model fitting criteria like AIC and BIC are presented in Tables 1–3. Effect sizes were estimated using G*Power 3.1.9.2. Reinstatement tests evoked by abbreviated ISS or non-contingent cue presentation were assessed using paired t-tests by comparing the responding on the active lever during reinstatement to the average of active lever presses on the last two extinction sessions. Reinstatement tests involved yohimbine were assessed using linear mixed-effects modeling followed by ANOVA.

**Table 1.** The statistical output from analyses testing the relation between behavioral markers and the demand for heroin.

|  | Model | Test | df | AIC | BIC | logLik | L.Ratio | p-value | Marginal R<sup>2</sup> |
| --- | --- | --- | --- | --- | --- | --- | --- | --- | --- |
| *Baseline Model (DV=EV)* | 1 |  | 3 | 91.16 | 93.48 | -42.58 |  |  |  |
| **Adding OF Center Diff** | **2** | **2 vs 1** | **4** | **84.54** | **87.63** | **-38.27** | **8.63** | **<0.01** | **0.43** |
| Adding OF Path Diff | 3 | 3 vs 1 | 4 | 91.98 | 95.07 | -41.99 | 1.18 | 0.28 | 0.07 |
| Adding OF Perimeter Diff | 4 | 4 vs 1 | 4 | 90.52 | 93.61 | -41.26 | 2.65 | 0.10 | 0.16 |
| **Adding FS Climbing** | **5** | **5 vs 1** | **4** | **82.40** | **85.49** | **-37.20** | **10.76** | **<0.01** | **0.50** |
| **Adding FS Immobility** | **6** | **6 vs 1** | **4** | **82.84** | **85.93** | **-37.42** | **10.32** | **<0.01** | **0.49** |
| Adding FS Swimming | 7 | 7 vs 1 | 4 | 92.97 | 96.06 | -42.49 | 0.19 | 0.66 | 0.01 |
| **Adding OF Center Diff and**<br>**FS Climbing together** | **8** | **8 vs 1** | **5** | **75.27** | **79.14** | **-32.63** | **19.88** | **<0.001** | **0.72** |

Significant effects are marked in **bold**.

**Table 2.** The statistical output from analyses assessing the relation between a combination of behavioral and biological markers and the demand for heroin.

|  | Model | Test | df | AIC | BIC | logLik | L.Ratio | p-value | Marginal R<sup>2</sup> |
| --- | --- | --- | --- | --- | --- | --- | --- | --- | --- |
| *Baseline Model (DV=Cort post ISS)* | 1 |  | 3 | 255.71 | 257.62 | -124.85 |  |  |  |
| **Adding Cort post FST** | **2** | **2 vs 1** | **4** | **251.52** | **254.07** | **-121.76** | **6.19** | **<0.01** | **0.37** |
| *Baseline Model (DV=EV)* | 1 |  | 3 | 81.78 | 83.7 | -37.89 |  |  |  |
| Adding Post-ISS Cort | 2 | 2 vs 1 | 4 | 90.49 | 93.58 | -41.25 | 2.67 | 0.10 | 0.16 |
| Adding Post-FST Cort | 3 | 3 vs 1 | 4 | 93.21 | 96.30 | -42.61 | 0.47 | 0.83 | 0.03 |
| **Adding Cort ISS/FST** | **4** | **4 vs 1** | **4** | **79.18** | **81.74** | **-35.59** | **4.6** | **0.03** | **0.29** |
| *Baseline Model (DV=EV)* | 1 |  | 3 | 81.78 | 83.7 | -37.89 |  |  |  |
| **Adding OF Center Diff**<br>**and Cort post ISS together** | **2** | **2 vs 1** | **5** | **74.64** | **77.84** | **-32.32** | **11.14** | **<0.01** | **0.56↑** |
| *Baseline Model (DV=EV)* | 1 |  | 3 | 81.78 | 83.7 | -37.89 |  |  |  |
| **Adding FS Climbing and**<br>**Cort post FST together** | **2** | **2 vs 1** | **5** | **70.73** | **73.93** | **-30.36** | **15.05** | **<0.001** | **0.67↑** |
| *Baseline Model (DV=EV)* | 1 |  | 3 | 81.78 | 83.7 | -37.89 |  |  |  |
| **Adding OF Center Diff,**<br>**FS Climbing, and**<br>**Cort ISS/FST together** | **2** | **2 vs 1** | **6** | **65.36** | **69.19** | **-26.68** | **22.43** | **<0.001** | **0.81↑** |

↑ - indicates an increase in R<sup>2</sup> when comparing to a behavioral predictor or predictors used in a corresponding model. Significant effects are marked in **bold**.

**Table 3.** The statistical output from tests assessing the relation between behavioral markers and the reinstatement behavior.

|  | Model | Test | df | AIC | BIC | logLik | L.Ratio | p-value | Marginal R<sup>2</sup> |
| --- | --- | --- | --- | --- | --- | --- | --- | --- | --- |
| *Baseline Model (DV=Cue)* | 1 |  | 3 | 149.83 | 152.14 | -71.91 |  |  |  |
| Adding Essential Value | 2 | 2 vs 1 | 4 | 149.71 | 152.80 | -70.85 | 2.12 | 0.15 | 0.13 |
| Adding OF Center Diff | 3 | 3 vs 1 | 4 | 151.15 | 154.24 | -71.57 | 0.68 | 0.41 | 0.04 |
| **Adding FS Climbing** | **4** | **4 vs 1** | **4** | **143.19** | **146.28** | **-67.59** | **8.64** | **<0.01** | **0.43** |

Significant effects are marked in **bold**.

## 4 Results

### 4.1 The effect of intermittent swim stress on social exploration and open field behaviors

Intermittent swim stress did not affect social exploration behavior (t(15)=0.42, p=0.68; Figure 2A). Intermittent swim stress significantly decreased time spent in the center of open field (t(15)=4.48, p<0.001), significantly increased time spent in the perimeter of open field (t(15)=4.85, p<0.001), and significantly decreased total distance traveled during the open field (pathlength; t=(15)=6.42, p<0.0001; Figure 2B-D).

**Figure 2.**
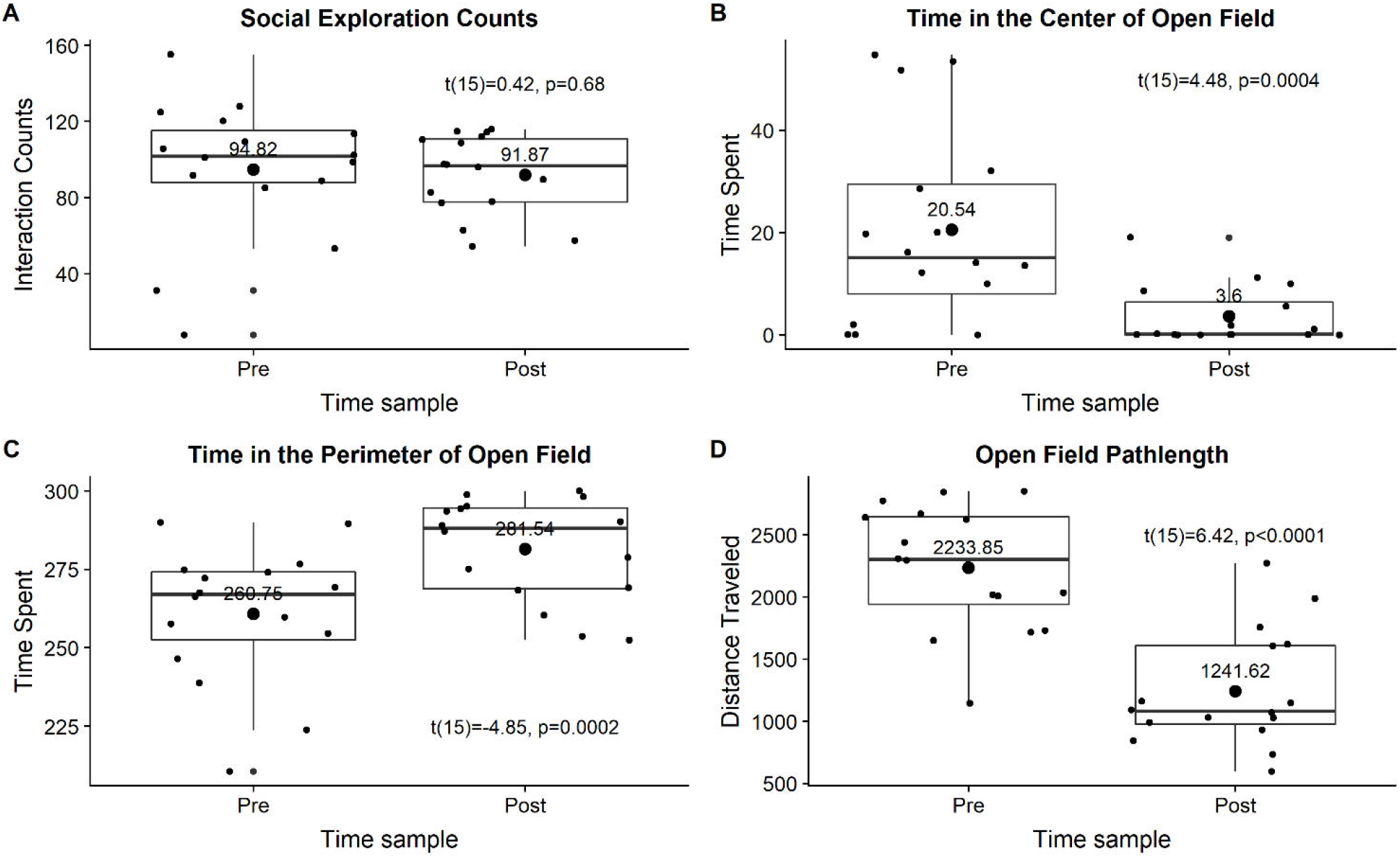
The effect of Intermittent Swim Stress on social exploration **(A)**, time spent in the center of the open field **(B)**, time spent in the perimeter of the open field **(C)**, or distance traveled during the open field test **(D)**. All data were visualized as grouped pretest and posttest measures using box plots with small points showing individual datum and large points showing means with mean values above.

### 4.2 Heroin self-administration and extinction

During the first 8 sessions of heroin self-administration responding on levers varied by Session (F(7, 240) = 10.13, p<0.0001), Lever (F(1, 240)= 212.6, p<0.0001), and there was no interaction (F(7, 240) = 1.205, p=0.30). Active lever responding was consistently higher over the first 8 sessions of heroin self-administration (Bonferroni comparisons; Figure 3A). The lowest individual demand (essential value) for heroin was 1.13 and the highest was 13.08 with the mean of 5.15, the standard deviation of 3.57, and coefficient of variation equal to 69.48% (Figure 3B). Active lever responding was consistently higher over the 5 sessions of self-administration following demand acquisition phase (F(1, 150)=409.7, p<0.0001; Bonferroni comparisons; Figure 3C). Lever responding over the 5 sessions of self-administration following demand acquisition phase did not varied by Session (F(4, 150) = 0.5653, p=0.68) and there was not Session by Lever interaction (F(4, 150) = 0.8616, p=0.4886). During extinction active lever responding decreased from 315.19 (SD=209) on session 1 to 44.44 (SD=35.72) on last session 10 (Figure 3D).

**Figure 3.**
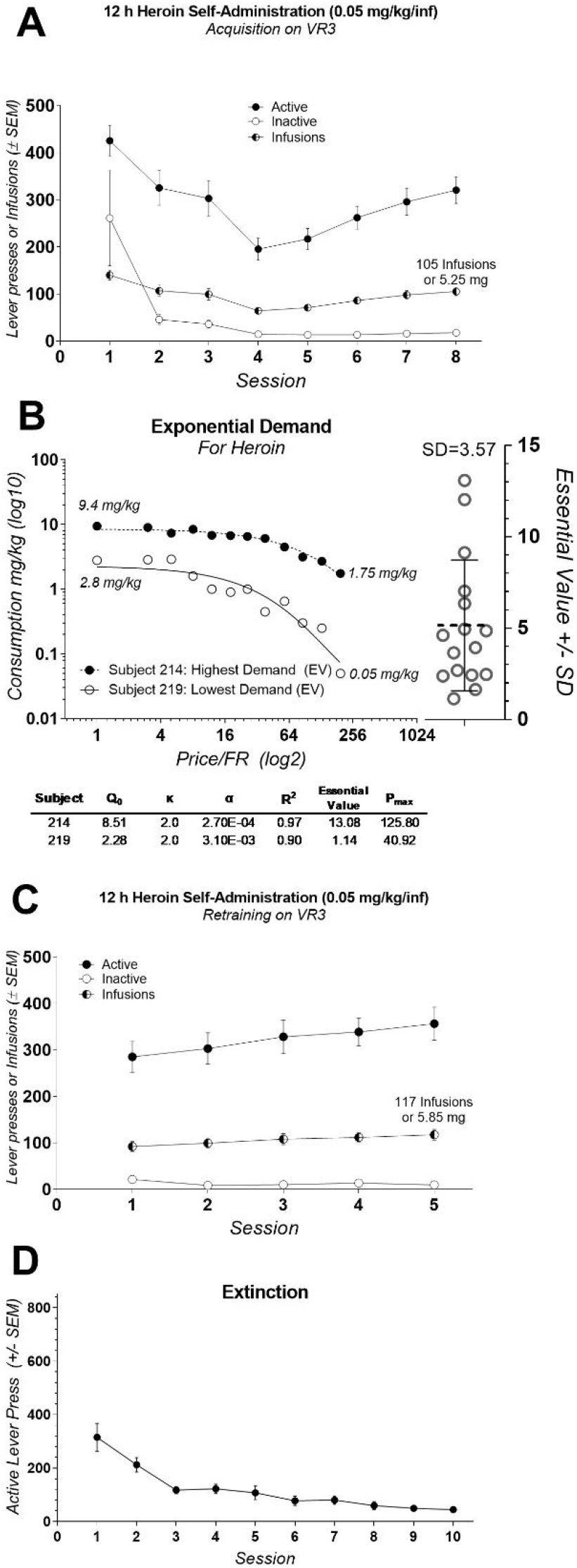
**(A)** Mean (±SEM) number of lever presses and infusions earned during the acquisition of heroin self-administration. **(B)** Exponential demand curves from rats with the highest and lowest essential value (left panel; critical demand values are shown below) and Essential Value scatterplot from all subjects (right panel). **(C)** Mean (±SEM) number of lever presses and infusions earned during reacquisition of heroin self-administration. **(D)** Mean (±SEM) number active lever presses during extinction.

### 4.3 Behavioral and biological markers predict demand for heroin

#### Individual behavioral markers are associated with increased demand for heroin

The detailed statistical output from the tests below is provided in Table 1. In the open field, neither the difference score in distance traveled nor the difference score in time spent in the perimeter of the open field before and after the ISS significantly related to the individual demand for heroin. Swimming during the FST did not correlate with the demand for heroin. On the other hand, time spent in the center of the open field after the stress minus the time spent in the center of the open field before the stress (difference score) significantly predicted individual demand for heroin (χ^2^(1)=8.63, p<0.01). The difference in the open field center activity explained approximately 43% of the variance in demand for heroin with the effect size (*f*^2^) equal to 0.75 (R^2^=0.43; Figure 4A). Climbing (χ^2^(1)=10.76, p<0.01) and Immobility (χ^2^(1)=10.32, p<0.01) during the 5 min FST, administered a day after the intermittent swim stress exposure, also significantly predicted an increased demand for heroin. Because climbing and immobility measures had an almost perfect negative relation with each other (R^2^ = 0.97), we are only showing and discussing climbing activity to avoid redundancy. Climbing activity during the FST explained approximately 50% of the variance in demand for heroin (R^2^ = 0.50; f^2^=1 Figure 4B). When both open field center difference score and the climbing during FST were added to the linear model at the same time, they improved the fit of the model by explaining a larger proportion of variance in demand for heroin then each predictor alone (χ^2^(2)=19.88, p<0.001; R^2^=0.72; f^2^=2.57). The fact that a combination of these measures improves the predictive ability of the linear model indicates that these measures explain a different portion of the variance in heroin demand and complement each other in their predictive qualities. Predicted values from the linear model that included open-field center difference score and climbing measure from FST as predictors for heroin demand are visualized in Figure 4C. Overall, these results indicate that difference in open field activity and climbing during the FST that followed the ISS are strong behavioral predictors of heroin demand that was assessed in a later phase of the study.

**Figure 4.**
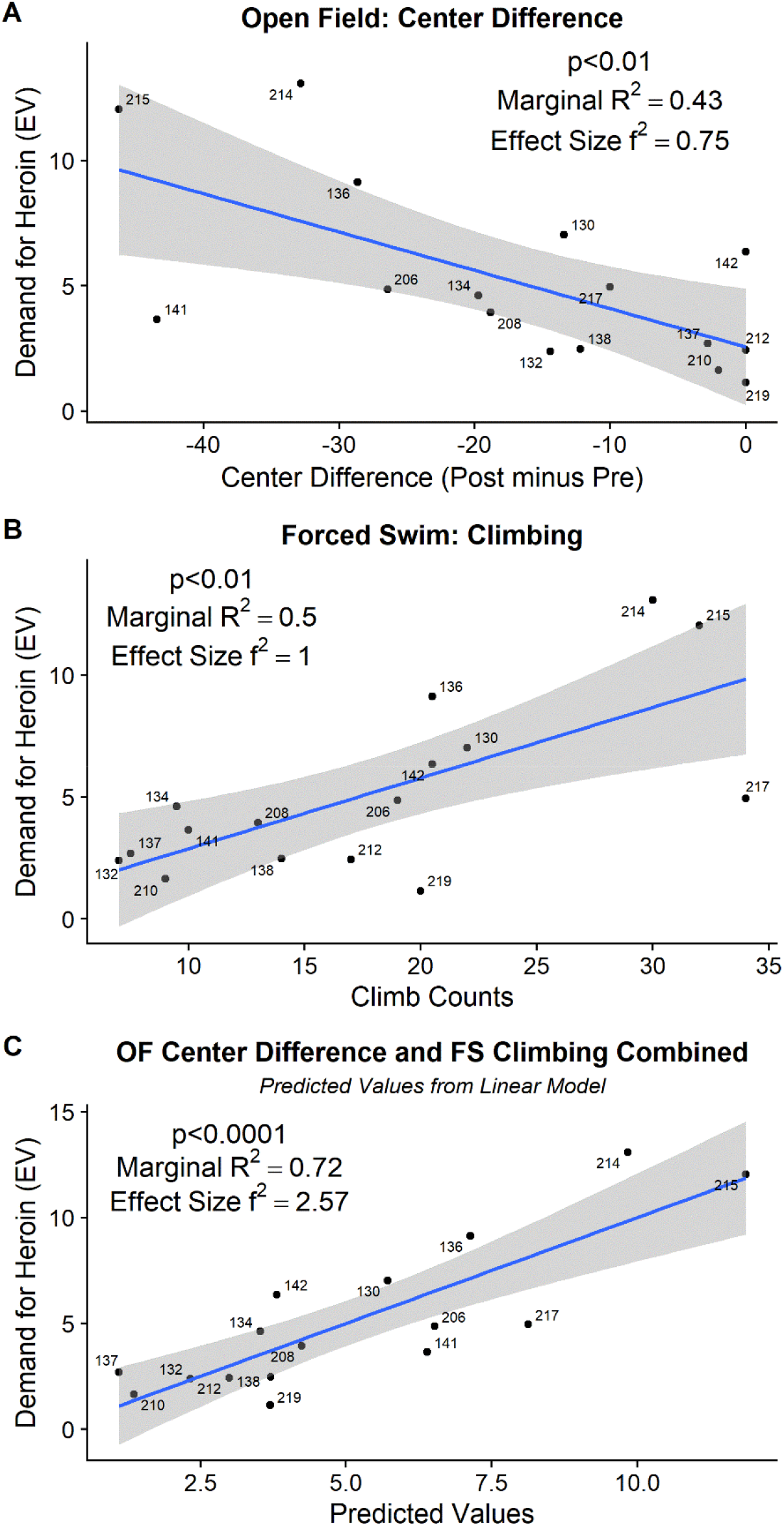
**(A)** The difference in time spent in the center of the open field, before and after the Intermittent Swim Stress, predicted subsequent demand for heroin. **(B)** Increased climbing during the 5 min of forced swim test, administered the day after Intermittent Swim Stress, predicted subsequent demand for heroin. **(C)** Together, open field difference score and forced swim climbing (visualized as predicted values from a linear model) explain 72% of the variance (R^2^) in subsequent demand for heroin.

#### Individual biological response to stress predicts demand for heroin

The detailed statistical output from the model fitting tests below is provided in Table 2. Plasma corticosterone levels were increased from baseline post-ISS and post-FST (F(3,12)=3.89, p < 0.05; Dunnett’s tests; Figure 5A). Individually, post-ISS or post-FST corticosterone responses did not predict demand for heroin; however, there was a positive linear relationship between these measures (χ^2^(1)=6.19, p<0.01; R^2^=0.37; f^2^=0.58; Figure 5B). To assess whether combined individual corticosterone response to ISS and FST related to the demand for heroin we created a corticosterone ISS/FST composite score by centering (z-score) both of these variables and summating them for each subject. With that in mind, combined corticosterone response to ISS and FST positively predicted demand for heroin (χ^2^(1)=4.60, p=0.032; R^2^=0.29; f^2^=0.41; Figure 5C) indicating that individual biological response to stress alone can be used to predict demand for heroin in the later phase of the study.

**Figure 5.**
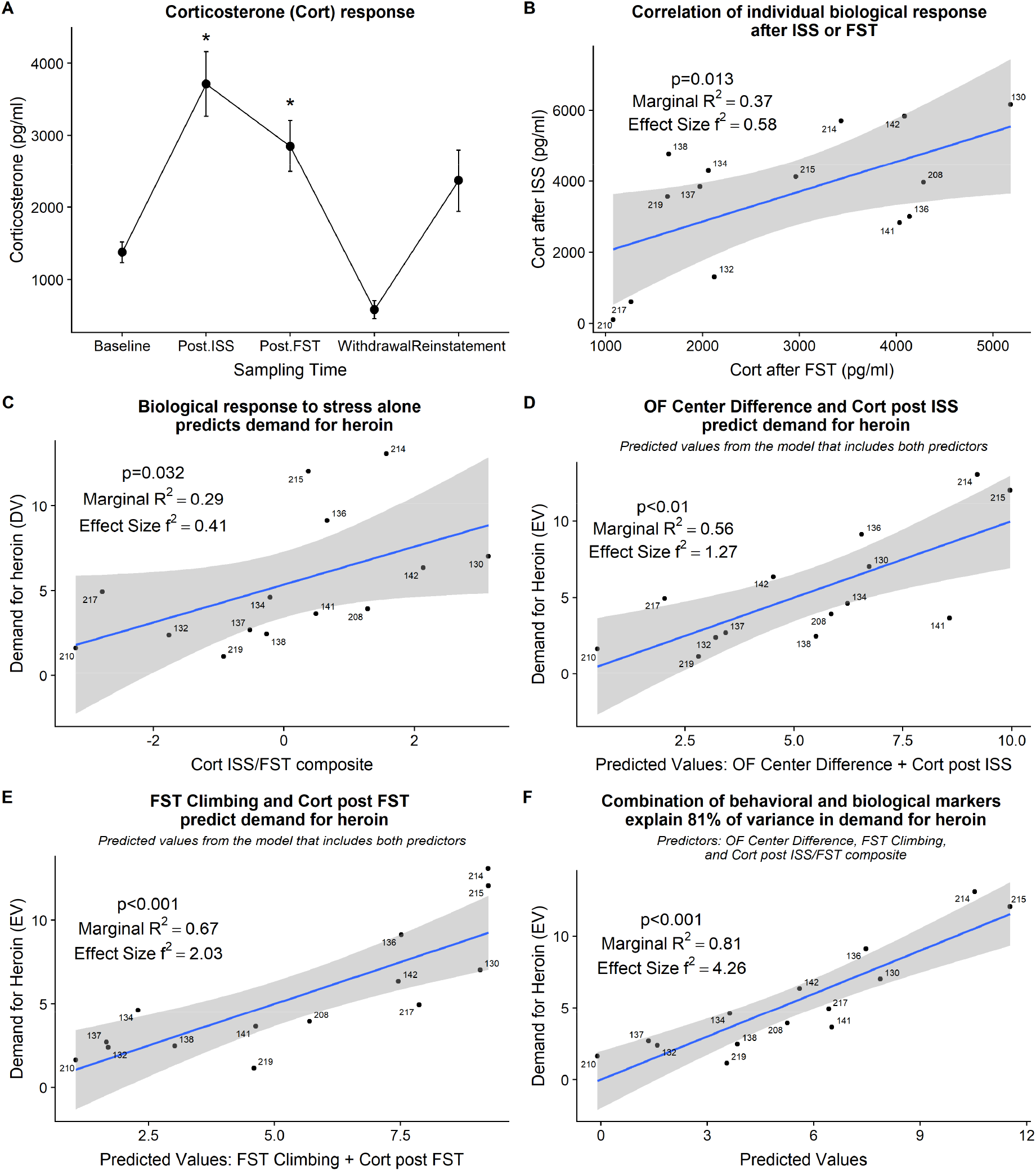
**(A)** Grouped corticosterone response at various points of the study. *Significant difference from baseline (p<0.05). *Post.ISS* – blood collected after ISS test. *Post.FST* – blood collected after FST. **(B)** Corticosterone response after ISS positively relates to corticosterone response after FST. **(C)** Corticosterone response after ISS and FST, combined into a composite score, predicts demand for heroin. **(D)** Individual open field center difference score and individual corticosterone response after ISS combined explain 56% of variance in demand for heroin. **(E)** Increased climbing during FST and increased corticosterone response after FST combined explain 67% of variance in demand for heroin. **(F)** Combination of behavioral and biological markers explain 81% of variance in demand for heroin. *ISS* – intermittent swim stress. *FST* – forced swim test. *OF* – open field. *Cort* – corticosterone.

#### Combination of behavioral and biological markers improves the ability to predict demand for heroin

We have shown above that behavioral and biological markers that were strategically sampled at time points relevant to a stress response induced by the ISS predict the demand for heroin in the later phase of the study. We then assessed how a combination of behavioral and corresponding biological markers relate to a demand for heroin. Open field center difference score and corticosterone response after the ISS together explained 55% of the variance in demand for heroin (χ^2^(2)=11.14, p<0.01; R^2^=0.56; f^2^=1.27; Figure 5D); R^2^ change = +0.13 when compared to a corresponding behavioral marker alone. Climbing during FST and corticosterone after the FST together explain 67% of the variance in demand for heroin (χ^2^(2)=15.05, p<0.001; R^2^=0.67; f^2^=2.03; Figure 5E); R^2^ change = +0.17 when compared to a corresponding behavioral marker alone. Finally, a combination of open field center difference score, climbing during the FST, and corticosterone ISS/FST composite together explain 81% of variance in demand for heroin (χ^2^(3)=22.42, p<0.001; R^2^=0.81; f^2^=4.26; Figure 5F); R^2^ change = +0.09 when compared to a model that includes both corresponding behavioral markers. These findings indicate that behavioral and biological markers are complimentary in their nature and together explain a large proportion of variance in individual demand for self-administered heroin.

### 4.4 Reinstatement

#### 4.4.1 Stress- and cue-triggered reinstatement

Abbreviated intermittent stress swim re-exposure (20 swim trials), administered before the 60 min extinction test, did not reinstate active lever responding when compared to the average of responding on the last two extinction sessions (t(15)=1.52, p=0.14; Figure 6A; compare ISS bar with Extinction bar). In contrast, non-contingent cue presentations during the extinction test significantly increased active lever responding when compared to the responding on the last two extinction sessions (t(15)=-3.68, p<0.01; Figure 6A; compare Cue bar with Extinction bar). Importantly, this is the first demonstration of heroin-associated cues presented non-contingently during extinction reinstating heroin seeking in rats with a history of stress exposure.

**Figure 6.**
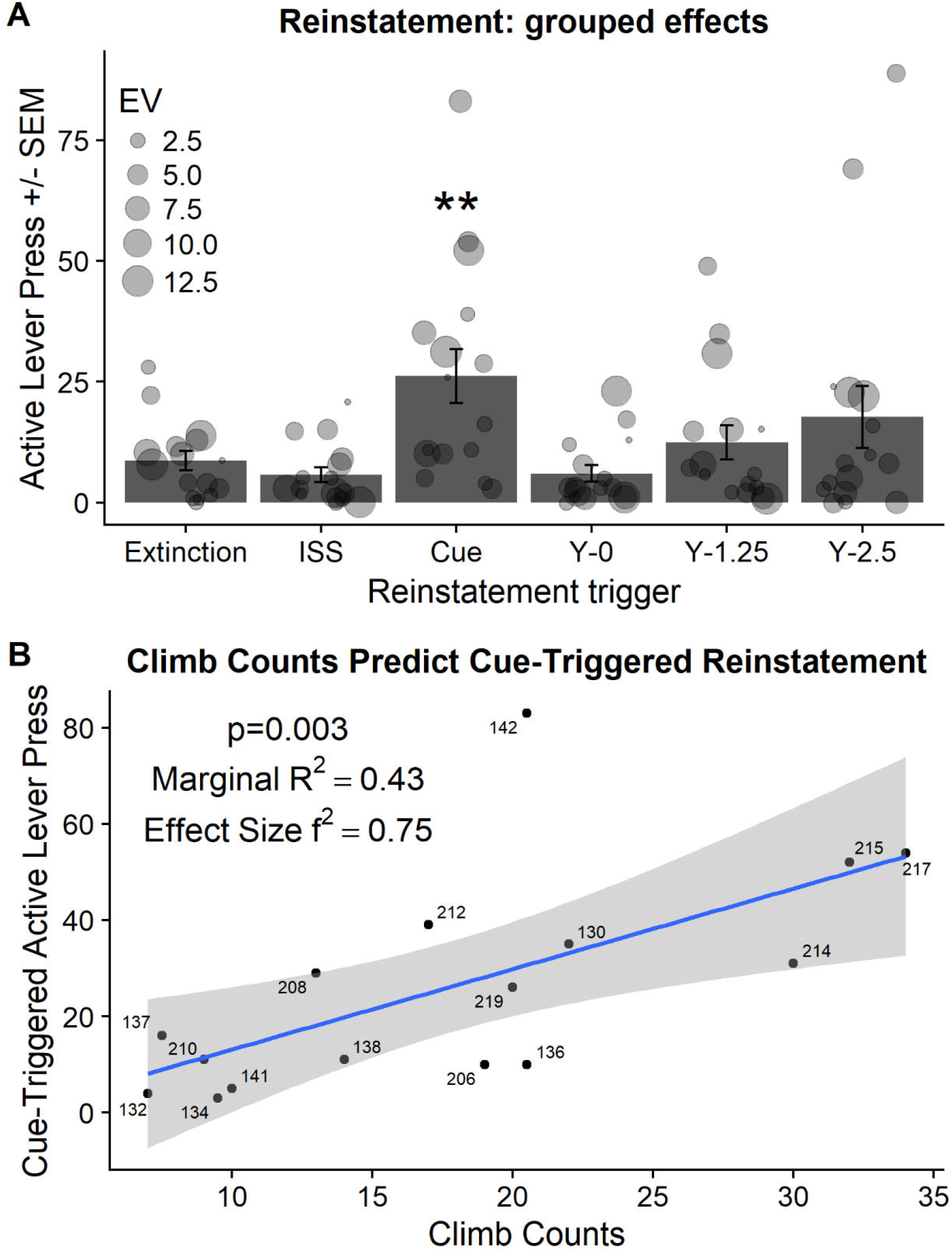
**(A)** Active lever presses (+/− SEM) during reinstatement tests compared to active lever presses in extinction (average of last two extinction sessions). EV-indicates the essential value or demand for heroin during self-administration phase of the experiment and is visualized using the size of the individual point. ISS – indicates abbreviated Intermittent Swim Stress that was used as a trigger. Y - indicates yohimbine pretreatment followed by the dose in mg/kg. **Significant difference from responding in extinction (p<0.01). **(B)** Active lever responding in extinction triggered by non-contingent cue presentations was predicted by climbing during the Forced Swim Test that was administered the day after the exposure to Intermittent Swim Stress.

#### 4.4.2 Yohimbine-triggered reinstatement

There was trending but not significant effect of yohimbine dose on active lever responding in extinction [F(2,30)=2.625; p=0.089; Figure 6A; compare bars labeled Y-0 to Y-2.5].

#### 4.4.3 Assessing the relationship between the demand for heroin, stress markers, and reinstatement behavior

Individual responding during non-contingent cue-triggered reinstatement was positively related to enhanced individual climbing during FST administered after the initial ISS exposure (χ^2^(1)=8.64, p<0.01; Figure 6B). Neither relevant behavioral markers (demand for heroin, open field center difference activity) nor relevant biological markers (corticosterone response to ISS/FST) predicted the magnitude of cue-triggered reinstatement.

## 5 Discussion

We hypothesized that individuals vulnerable to the effects of stress have a higher risk of abusing heroin after the exposure to stress. We further hypothesized that we could simulate these individual effects in a preclinical model of stress and heroin use comorbidity. In this model, all rats are initially exposed to a stress episode and then their individual biological response to stress in a form of corticosterone response and individual behavioral responses that are strategically sampled around the stress exposure are used to predict individual levels of heroin self-administration that are derived using a behavioral economics model. Importantly, in this experimental design the time between the stress exposure and the initiation of heroin self-administration models clinically relevant time interval between stress and drug use. Using this approach, we show that a) corticosterone response after ISS and FST was higher than at baseline, b) the magnitude of the corticosterone response after ISS and FST combined predicted the magnitude of the demand for heroin, c) behavioral markers strategically sampled around the time of stress exposure predicted the demand for heroin, and d) a combination of biological and behavioral markers greatly improved the ability to predict individual demand for heroin, when compared to biological or behavioral markers alone. Furthermore, for the first time, we show that heroin-associated cues, which were non-contingently presented during extinction, reinstated heroin seeking in extinction in rats with a history of stress exposure. Importantly, higher climb counts from FST, that was administered the day after ISS, predicted a higher magnitude of cue-triggered reinstatement. Overall, these findings show that the individual biological response to stress and individual behaviors sampled around the time of stress exposure can be used as predictors for the magnitude of stress-induced heroin self-administration and cue-triggered reinstatement of drug seeking in extinction.

The protocol employed in the current study was designed to simulate progression from stress exposure to stress-induced opioid use. To induce long-lasting effects of stress, with a high degree of ecological validity, we used the ISS paradigm (Brown et al. 2001; Christianson and Drugan 2005; Drugan et al. 2014; Stafford et al. 2015). The ISS paradigm was developed as a hybrid model combining the strengths from two other animal models: 1) the use of the ecologically valid stressor of inescapable swim stress (Porsolt et al. 1977), and 2) the unpredictable and inescapable intermittent stress exposure from the learned helplessness paradigm (Maier and Seligman 1976). The severity of the stressor is furthermore controlled a priori by the experimenter so that each subject receives the identical amount of stress and therefore allows direct comparison across groups or individual subjects (Christianson and Drugan 2005; Drugan et al. 2005, 2014; Stafford et al. 2015). In this study, we show that ISS evoked significantly higher corticosterone response when compared to the no-stress baseline. This finding is consistent with our previous results demonstrating that the ISS in 20 or 25 °C water evokes comparable serum levels of corticosterone (~3500-4000 pg/ml; Drugan et al. 2005). Although individual corticosterone response after the ISS did not predict the demand for heroin in our study, possibly demonstrating the lack of statistical power, we do show that the magnitude of combined corticosterone response after the ISS and the FST positively related to a demand for heroin assessed weeks after the stress episode. Importantly, the corticosterone response after the FST was also significantly higher than no-stress baseline levels and was positively correlated with the corticosterone response after the ISS. This positive relationship between the biological response to these two swim stressors indicates that rats that are vulnerable to the effects of ISS also exhibit higher stress response after the FST that was administered the day after the ISS and was designed to assess the short-term effects of ISS. These results indicate that biological response to stress alone in the form of serum levels of corticosterone can be used to predict subsequent demand for heroin and that a combination of ISS and FST may serve as a reliable model to study individual long-term effects of stress and its interaction with other pathologies (e.g., substance use disorder).

To assess various modalities of stress-induced deficits on the individual level and relate them to heroin self-administration behavior, we administered three widely validated tests that model anxiety and helpless behaviors. Rats underwent social exploration and open field tests 24 hours before ISS and the same tests followed by a 5 min FST 24 hours after stress exposure. We found that greater decrease in time spent in the center of the open field and greater bouts of climbing during the FST predicted higher demand for heroin. Reduced time spent in the center area of the open field is widely validated as an analog of anxiety and stress induces long-term changes in this behavior (van Dijken et al. 1992; Prut and Belzung 2003; Hale et al. 2008). Likewise, the FST has been widely used to assess the effects of stress in preclinical models including the effects of ISS. For example, exposure to ISS has been shown to increase immobility in a subsequent FST in comparison to non-stressed controls, however, the individual variability has not been previously assessed (Christianson and Drugan 2005; Drugan et al. 2010). Interestingly, our study shows that the increased climbing during FST that was administered the day after ISS predicted greater heroin demand. Forced swim climbing behavior is usually decreased following prior swim exposure in the typical 2-day behavioral despair paradigm (Detke et al. 1995; Drugan et al. 2010). However, the interpretation of forced swim behaviors has been debated for decades (Nishimura et al. 1988; Commons et al. 2017). The alternative argument contends that passive behaviors during subsequent swim exposures are an adaptive coping strategy, whereas climbing movements may reflect susceptibility (Commons et al. 2017). It is important to note that our FST followed an initial stress experience with water as an aversive stimulus; by that time rats likely developed conditioned response to water submersion – conditioned avoidance. Also, hyperarousal or hyperreactivity is commonly associated with long-term effects of stress (Pynoos et al. 1996; Strekalova et al. 2004; Pibiri et al. 2008; Schöner et al. 2017). Our statistical analysis shows that the elevated climbing is a complementary measure to the deficits in open field activity and both measures significantly improve the predictive ability of the mixed-effects linear model to explain variability in demand for heroin. Also, adding climb counts during the FST and corticosterone response after the FST together into one predictive model significantly improved the ability to predict the demand for heroin when compared to each of those predictors alone. With all this in mind, we argue that elevated climbing during the FST test, which followed a stress episode, likely represents “conditioned hyperreactivity” and is consistent with signs of stress vulnerability.

To closely approximate human drug-taking condition and to study relevant behavioral and neurobiological processes we used a long-access (12 hours) preclinical model of drug self-administration in conjunction with a well-validated economic demand approach to study individual differences in stress-induced heroin drug taking. Early preclinical models of drug self-administration used unlimited access, often resulting in overdose-related deaths (e.g., Johanson et al. 1976), prompting adaptation of short-access protocols that are more economic, allow higher throughput, but lacking translational relevance (for more see Gawin and Kleber 1988; Gawin 1991). Importantly, using this protocol and VR_3_ schedule of reinforcement, rats self-administer large amounts of heroin per session (5.25 mg/kg during 8^th^ self-administration session; 0.05 mg/kg/inf) when compared to previous reports. For example, Kenny et al., (2006) used 23 h access protocol (0.02 mg/kg/inf) on fixed ratio schedule of reinforcement (FR1) and showed total consumption after first 8 sessions of heroin self-administration to be approximately 0.8 mg/kg and after 24 consecutive sessions to be approximately 1.8 mg/kg. In addition, Lynch and Carroll (1999) used 6 h access (0.015 mg/kg/inf) on FR1 and showed total consumption after 5 sessions to be approximately 0.75 mg/kg. Thus, using our 12 h access approach and a variable schedule of reinforcement we are able to ensure high heroin intake, robust lever discrimination, and no overdose-related deaths. Using this approach, we show that individual reactivity to stress was a strong predictor for increased heroin demand. Because this is a first study to show these individual effects in male rats, additional studies will be required to further confirm and extend these findings.

Relapse is a critical factor contributing to substance use and abuse (Hendershot et al. 2011). Clinical and preclinical studies show that relapse can be precipitated by stress and cues that have been previously associated with the drug (Sinha et al. 2011; Mantsch et al. 2016). For example, intermittent footshock, or pharmacological stressors like metyrapone or yohimbine, trigger higher rates of heroin seeking in extinction (Shaham and Stewart 1995; Shaham et al. 1996, 1997, 1998; Banna et al. 2010). Cues that have been previously paired with heroin infusions and that are presented contingently during extinction, those that require a response to earn a cue presentation, also reliably reinstate drug seeking in extinction (Banna et al. 2010; Doherty and Frantz 2012). In comparison, previous studies show that non-contingent cue presentations during extinction, that is when cues are presented by the experimenter and do not require a response, do not reinstate cocaine or heroin seeking (Alderson et al. 2000; Grimm et al. 2000). In contrast with the previous reports (Banna et al. 2010), we show that yohimbine, a pharmacological stressor with high affinity for the *α*_2_-adrenergic receptor that induces noradrenergic tone, did not reinstate heroin seeking in rats with the previous history of stress although the grouped effect was trending towards significance (p=0.089). Importantly, the variance observed at the highest doses indicates that some rats had a much greater response to yohimbine that others (observe individual response evoked by yohimbine that is visualized in Figure 6A). In addition, we show that brief ISS exposure also did not reinstate heroin seeking. This lack of effect following brief ISS exposure may be partially explained by the temporal dissociation from the reinstatement test, which was administered 30 min later, or by the limited stress effects induced by 20 trials rather than the prolonged 100 trials administered during the initial stress episode. Finally, one of the most interesting findings in our study is the reinstatement of heroin seeking by non-contingent cue presentations during extinction in rats with a history of stress exposure. In our study, non-contingent cue presentations consisted of lever retractions and illumination of both cue lights for 20 seconds, a sequence that was repeated every five minutes from the beginning of the session. The magnitude of cue-induced responding was predicted by the climbing behavior during the FST administered the day after the initial ISS stress induction.

Clinical and preclinical studies show that stress is an important factor in substance use disorders but there is also evidence of individual variability across various phases of stress and substance use comorbidity. These individual effects involved in stress and substance use comorbidity are not well studied and presently are not well understood. To advance towards more effective and individualized prevention and treatment strategies targeting substance use disorder there is a need to better understand how individual vulnerability or resilience to stress interacts with drug use. The study presented here starts filling this gap by outlining a framework for predicting individual economic demand for self-administered heroin based on biological and behavioral markers strategically sampled around the stress episode. To this end, our findings show that behavioral, biological, and a combination of behavioral and biological markers sampled prior and after the stress episode that occurred weeks before the access to heroin self-administration can predict individual demand for heroin. We also show that the individual biological response to stress can be measured by assessing individual corticosterone response evoked either by ISS or FST and then this biological response can be used to predict subsequent demand for heroin. For example, rats with higher corticosterone response to stress and higher demand for heroin can be conceptualized as vulnerable to stress and heroin use comorbidity phenotype. On the other hand, it is unclear that the behavioral responses following the stress episode were directly affected by the exposure to stress. Because our study does not include a no-stress control condition required to make such an assessment, we are not able to make a claim that a change in behavioral responses after the ISS is stress induced. However, because we show that behavioral responses sampled before and after the stress episode complement biological markers in their ability to predict the demand for heroin it is likely that they are related to the effects of stress although additional studies with appropriate controls are necessary to confirm this speculation. With all this in mind, our demonstration that a combination of biological stress markers and behavioral responses sampled before and after the stress episode can explain most variance (81%) in subsequent demand for heroin provides a framework for a variety of future studies that can further investigate behavioral, biological, or mediating factors underlying this effect.

## 6 Acknowledgments

Funding: S. Charntikov was partially supported by GM113131 (CIBBR, P20) while preparing this manuscript for publication. Partial funding was provided by the Undergraduate Research Opportunities Program at the University of New Hampshire awarded to C. Donovan and E. Hart. The authors would like to thank Monica Ford Ortiz, Daniel Hertia, and Kathryn Taylor for their assistance with behavioral procedures.

